# Experience-dependent refinement of natural approach responses towards specific visual stimuli in mice

**DOI:** 10.1101/2020.05.14.096941

**Authors:** Nicole M. Procacci, Kelsey M. Allen, Gael E. Robb, Rebecca Ijekah, Jennifer L. Hoy

## Abstract

Specific features of visual objects innately draw orienting and approach responses in animals, and provide natural signals of potential reward. In addition, the rapid refinement of innate approach responses enhances the ability of an animal to effectively and conditionally forage, capture prey or initiate a rewarding social experience. However, the neural mechanisms underlying how the brain encodes naturally appetitive stimuli and conditionally transforms stimuli into approach behavior remain unclear. As a first step towards this goal, we have developed a behavioral assay to quantify innate, visually-evoked approach behaviors in freely moving mice presented with simple, computer generated stimuli of varying sizes and speeds in the lower visual field. We found that specific combinations of stimulus features selectively evoked innate approach versus freezing behavioral responses. Surprisingly, we also discovered that prey capture experience selectively modified a range of visually-guided appetitive behaviors, including increasing the probability of approach and pursuit of moving stimuli, as well as altering those visual features that evoked approach. These findings will enable the use of sophisticated genetic strategies to uncover novel neural mechanisms underlying predictive coding, innate behavioral choice, and flexible, state-dependent processing of stimuli in the mouse visual system.

**Highlights:**

- Novel stimuli with specific visual features reliably elicit an approach in C57BL/6J mice.
- Introduction of motion to stimuli makes freezing the most probable behavioral response.
- Spontaneous behavioral responses are tuned to size, speed and visual field location.
- Prey capture experience selectively refines natural, visually-evoked approach behaviors.

## Introduction

The ability to rapidly orient towards prey or away from predators from a distance is crucial for the survival of animals. Thus, visual systems are evolutionarily honed to extract the specific visual cues that reliably distinguish objects as potentially rewarding or threatening and rapidly drive appropriate orienting behaviors (Dean, Redgrave, and Westby 1989; Knudsen 2020). Particular sizes, speeds of motion, and relative location in the environment are often characteristics of natural predators or prey. Therefore, specific combinations of these features are highly salient and robustly drive rapid visual orienting responses across the animal kingdom. For example, classical studies in the toad have revealed that specific shapes, orientations, and speeds of stimuli robustly release predatory striking behaviors versus those that release active avoidance (Ingle 1971; Ewert 1980). Similarly, specific sizes and speeds of two-dimensional visual stimuli evoke rapid orienting towards or away from moving stimuli in the larval zebrafish (Bianco, Kampff, and Engert 2011; Barker and Baier 2015). Despite the ability of specific combinations of visual features to drive robust orienting responses across the animal kingdom, it would be costly to release these energetic or risky behaviors in the wrong context. For example, even in the presence of a potentially appetitive stimulus that should drive approach behavior, animals should suppress this response when sated or in the presence of a competing, more salient cue that indicates possible threat (Lima and Dill 1990; Burnett et al. 2016). Thus, innate approach responses must be triggered by specific combinations of visual features that indicate reward, but they must also be flexibly modulated as a function of internal state, context, or experience on many distinct timescales (Filosa et al. 2016). Currently, the neural mechanisms underlying conserved visually-guided approach responses in mammals and their context-dependent modulation remain unclear.

The mouse has emerged as a powerful model to determine the circuit mechanisms underlying visually driven orienting behaviors such as approach, as well as the decision-making processes underlying choice of approach versus avoidance behaviors, or even choice between competing stimuli of similar valence. Prior studies in the mouse have shown that specific features of stimulus motion and their location in the upper visual field reliably drive selection of freezing versus fleeing behaviors (Yilmaz and Meister 2013; De Franceschi et al. 2016). These robust behavioral responses are highly conserved across species, indicating that the visual field location where stimuli appear conveys valence information and may ultimately be dissected to reveal the circuit mechanisms that mediate response choice to potential threats. On the other hand, recent work has begun to reveal the neural circuit basis of visual stimulus selection to conditioned visual stimuli that predict food or water rewards in head-restrained mice (Krauzlis et al. 2020; Lutas et al. 2019). However, few studies have quantified the features of novel visual stimuli that cause mice to spontaneously approach. Doing so would allow us to identify the neural activity that innately encodes appetitive stimuli and determine how it changes with experience or under distinct contexts. Under ecologically-relevant conditions, salient visual stimuli detected near the horizon may be appetitive or aversive, and mice ultimately have to choose quickly which stimuli to approach and which to avoid. For example, small, moving objects in this location might indicate potential prey, as is often cited (Dean, Redgrave, and Westby 1989). Conversely, similar simple cues could indicate social or predatory threat. Understanding how mice naturally respond to specific features of novel visual information in this region of the environment and select appropriate behavioral responses is therefore an important first step towards significantly expanding the utility of the mouse to understand the neural basis of visual perception, object encoding, decision making and modulation of selective attention.

Here, we present mice with virtual stimuli near the horizon of the environment that reliably evoke approach. While we find that mice approach a range of novel object sizes and speeds of motion, they show a clear preference to approach objects that are a specific size and speed. In addition, mice are able to accurately intercept moving stimuli without the benefit of explicit experience with the stimulus. Surprisingly, we find that mice frequently exhibit prolonged periods of immobility, or freezing, in response to our visual stimuli, and that this behavioral response constitutes a majority of stimulus-evoked responses. These freezing responses precede about 50% of accurate and successful approaches towards novel moving objects and thus they are evoked in a context distinct from that created by the presentation of overhead looming or sweeping moving stimuli. Most intriguingly, prey capture experience prior to exposure to novel moving stimuli robustly increases the ratio of approach to freezing responses by selectively altering those stimulus features that drive approach. Taken together, these findings support the idea that approach and freezing constitute two classes of related, yet distinct orienting responses innately evoked by specific combinations of stimulus size, speed, and location in the visual field. Moreover, approaches are flexibly and selectively modulated by prey capture experience, again suggesting that they are dissociable from freezing responses. This new understanding of how mice innately respond to specific combinations of visual features viewed near the horizon, and how prey capture experience selectively modifies these visual responses, will facilitate studies that identify novel neural mechanisms underlying predictive coding, object salience and valence, selective attention, decision making, and motivation.

## Results

### Novel, computer generated visual stimuli reliably elicit approach in C57BL/6J mice

Previous work has shown that mice use visual cues to recognize prey (Hoy et al. 2016; Hoy, Bishop, and Niell 2019; Shang et al. 2019; Zhao et al. 2019). In other species, specific combinations of visual features innately elicit approaches towards possible prey items (Barker and Baier 2015; Bianco, Kampff, and Engert 2011; Ingle 1971). However, the specific visual features of prey that draw innate approach responses in mice are unclear. In order to parametrically vary the visual stimulus to determine the features that drive approach behavior in mice, we developed an assay employing computer generated stimuli projected onto a computer screen to elicit innate orienting responses in freely-moving mice. We parametrically varied the size (**Fig. 1**) or speed (**Fig. 2**) of computer generated, black ellipses on a white background. First, a single cohort of mice was presented with a stationary, black ellipse on one of two computer screens comprising two of the four sides of an open field arena (**Fig. 1A**, blue outlines). The ellipse was centered on one of three possible locations along the azimuth of the target screen and the bottom edge was maintained at 1 cm from the floor in elevation. The stimulus had a horizontal axis twice as long as the vertical axis in order to model potential insect prey at this location in the environment. Maintaining this aspect ratio, we randomly varied the size of the stimulus along the horizontal axis from 0.5 to 8 cm, and quantified behaviors elicited within 60 s of the start of stimulus presentation. To derive behavioral measures, we tracked the stimulus as well as the nose, ears, and tail base of the mice (**Fig. 1B**). We calculated the mouse’s approach frequency, range (distance between stimulus and mouse head at approach start), locomotion speed, and head angle relative to stimulus. Importantly, studies have shown that eye movements are coupled to head position in space and are aligned to head direction when performing natural visual behaviors such as prey capture and social investigation (Meyer, O’Keefe, and Poort, n.d.; Michaiel, Abe, and Niell, n.d.). Thus, head angle is a reliable measure of probable viewing angle and likely visual gaze. We defined a successful approach as any time the mouse’s nose came within 2 cm of the stimulus center. Approaches were identified as in Hoy et al., 2016. Briefly, an approach start was defined as when mice decreased both their range and head angle relative to the stimulus while moving an average speed of at least 15 cm/s starting from at least 5 cm away (**Fig. 1C**). Mice almost completely failed to approach stimuli 0.5 cm in size along the horizontal axis (**Fig. 1D-F**), and significantly slowed approach speeds to stimuli larger than 2 cm (**Fig. 1E**). In contrast, mice approached the 2 cm diameter stimulus most frequently, and with the highest locomotive speeds (**Fig. 1D & E**). Thus, while mice approach a range of stimulus sizes, they preferred stimuli that were 2 cm in length along the horizontal axis. Moreover, we observed that approach starts systematically occurred from farther ranges as stimulus size increased (**Fig. 1F**). Given the nearly linear relationship between the distance where an approach started and stimulus size (**Fig**.**1F**), we estimated the preferred relative stimulus size as the slope of a linear fit to the data. This yielded a preferred relative stimulus size of approximately 5 degrees (deg) of the visual angle. We were therefore able to determine that mice spontaneously orient towards and approach a preferred size of stimulus, as predicted from egocentric head position.

**Figure 1.**
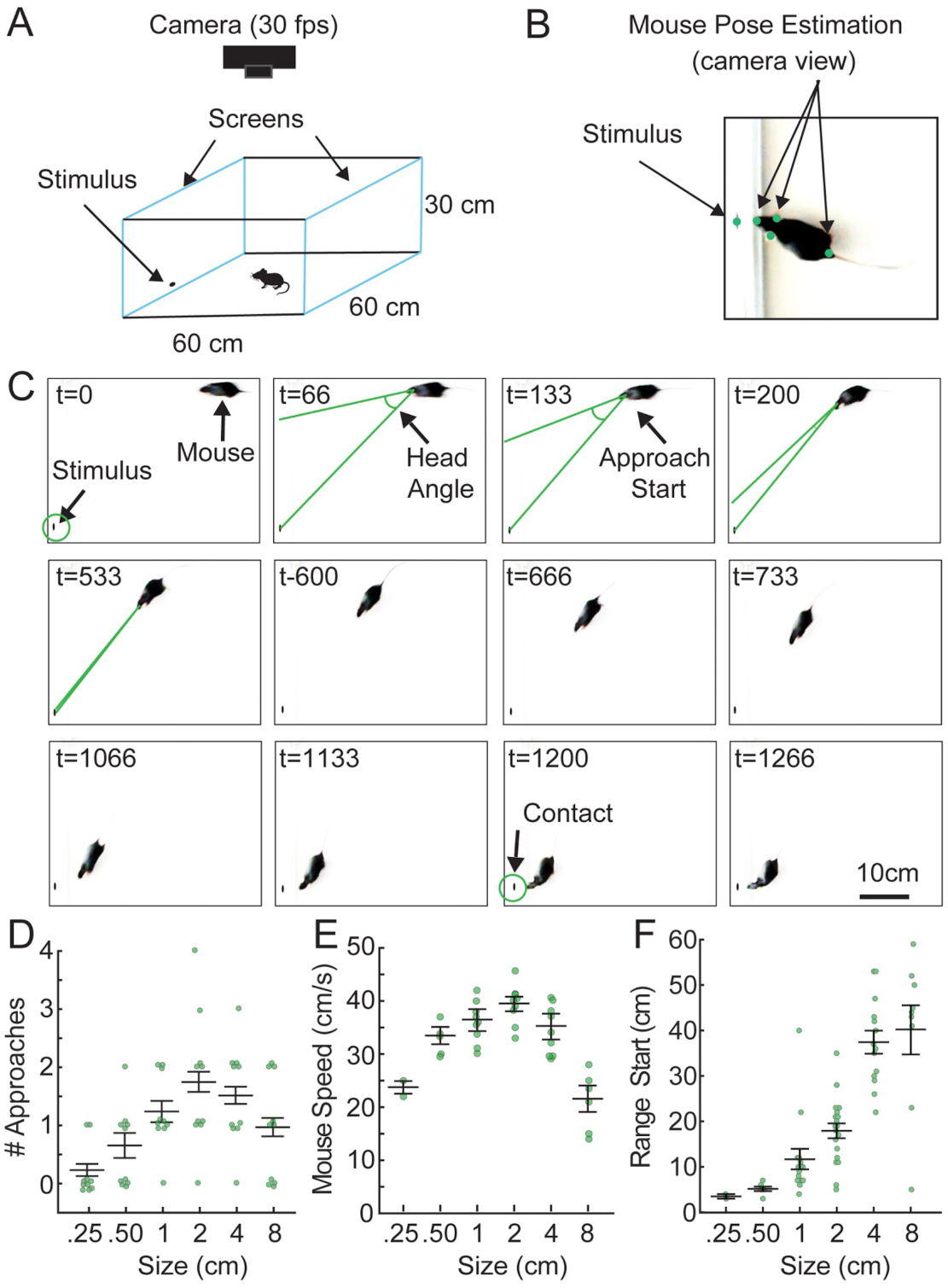
Naive mice preferably approach specific sizes of stationary, two-dimensional visual stimuli. Naive mice have only been habituated to the open field and handlers, but have not been exposed to computer generated stimuli or live crickets previously. (A) Schematic of behavioral arena with video recording performed overhead. Two sides of a white acrylic behavior box consisted of computer monitors displaying blank white screens. For experiments with stationary stimuli, ellipses of different sizes were presented in different locations within 1 cm of the bottom of one of the monitors in randomized orders. (B) Example frame from a recorded behavioral video. Egocentric positional information (range and head angle) between the mouse and stimulus are calculated from pose estimation data (see also **Video 1** employing moving stimul**i**). The stimulus position is tracked relative to the mouse’s nose, ears and tail base shown as green circles in the schematic. (C) A representative approach sequence displayed by a mouse towards a stationary ellipse presented on one of the monitors, highlighting specific moments surrounding a successful approach: approach start and contact. Green lines indicate mouse head angle relative to stimulus center. (D) Mean approach frequency for each mouse exposed to 6 different objective sizes of stimuli. N = 10 mice. (E) Mean locomotor speed for mice that approached stimuli. N = 2, 5, 9, 9, 9 and 6 mice each stimulus, respectively. (F) Mean range of approach starts at each objective size of stimulus. N = 65, r = 0.73, adjusted r^2^ = 0.487. Error bars are +/- standard error of the mean (SEM) in all panels.

**Figure 2.**
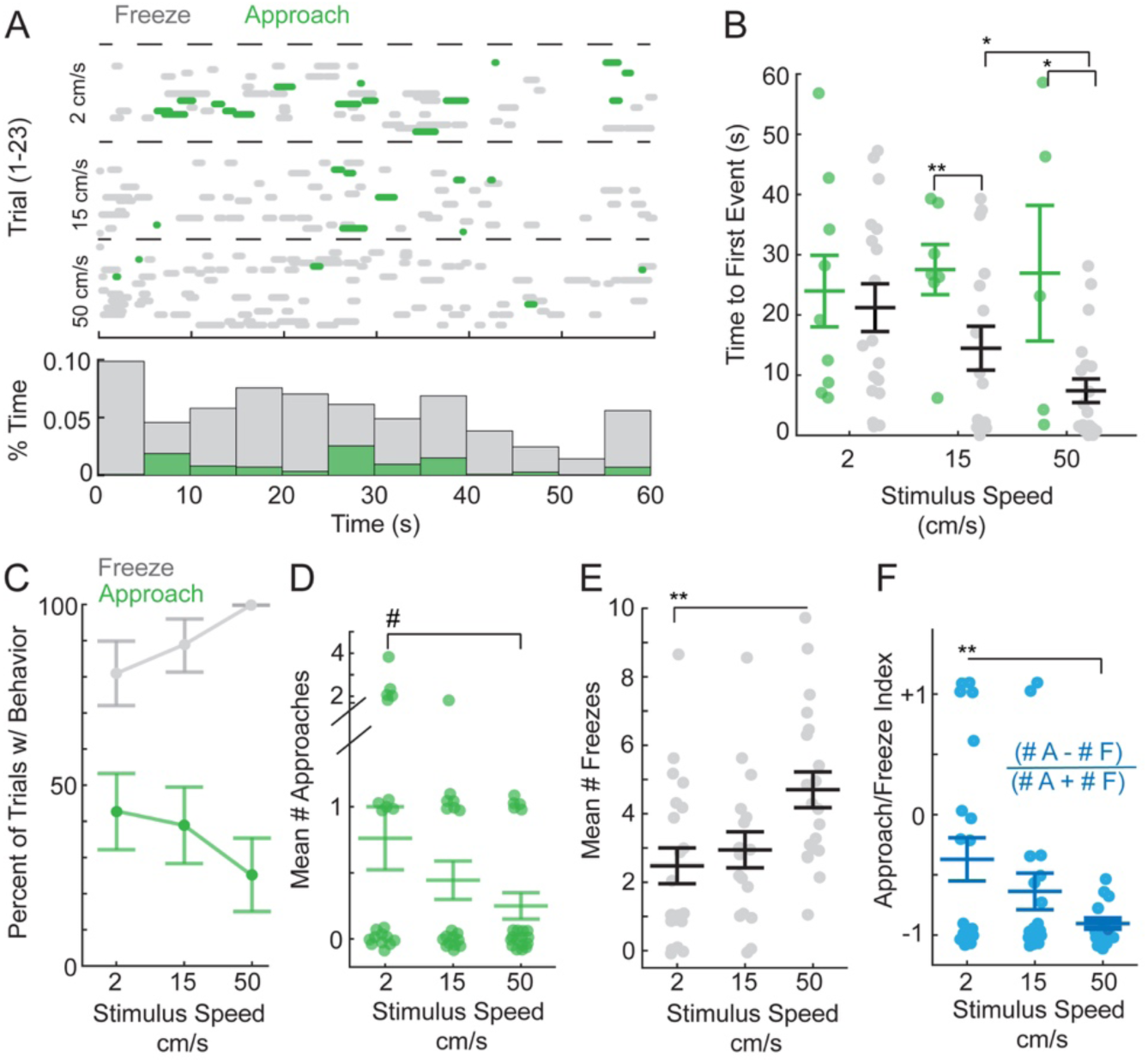
Mice freeze and/or approach when presented with novel, small, moving stimuli. (A) Top, an ethogram representing when approach or freezing (period of immobility > 500 ms) occurred from the onset of presentation of three different speeds of the stimulus for each mouse (each trial is denoted by a separate line in the ethogram). Green = approaches, grey = freezes. Dashed line separates trial types based on stimulus speed, N = 23 mice performing one trial at each of three stimulus speeds. Bottom histogram, the proportion of time spent engaged in each type of response during each 5 s segment of the 60 s trial, collapsed across all speeds. (B) Mean time to the first behavioral event of each type for mice that displayed at least one type of orienting response, green = approaches and grey = freezes (two-way ANOVA, F= 9.1957, p < 0.001, Tukey’s post hoc, * = p < 0.05 and ** = p< 0.01, Ns for each speed = 9, 7, 5 mice and 17, 16, 20 mice exhibiting approaches versus freezing, respectively). (C) Percentage of trials at each speed with at least one freeze (grey) or approach (green). Fisher’s exact tests indicated the percentage of trials with freezes was significantly higher than that with approaches for each stimulus speed (p < 0.05, p < 0.01 and p < 0.0001, respectively). Error bars are standard deviation. (D) Number of approaches per mouse (one-way ANOVA, (F(2) = 2.8418, p = 0.07, # = nearly significant). All conditions had significantly more approaches than a hypothetical mean of 0 (Student’s t-tests, p < 0.001, p < 0.01 and p < 0.05). (E) Number of freezes per mouse (one-way ANOVA, (F(2) = 6.54, p < 0.01, Tukey’s post hoc, ** = p < 0.01). Freezing evoked by all stimuli were highly significantly greater than a hypothetical mean of 0 (Student’s t-tests, p < 0.0001 in all cases). (F) Approach-to-freeze index calculated as shown in order to determine how relative response probability changes by stimulus speed (one-way ANOVA, (F(2) = 5.97, p < 0.01, Tukey’s post hoc, ** = p < 0.01). Error bars are +/-SEM.

### Stimulus speed biases the probability of approach versus freezing behavior

Given that dynamic stimuli are biologically salient and evoke a range of innate orienting behaviors, we next measured the behavioral responses of freely-moving mice to stimuli that moved along the azimuth with steady speeds. The stationary stimulus that evoked the most approaches was 2 x 1 cm, thus we varied the speed of this stimulus, ranging from 2 cm/s to 50 cm/s in order to determine whether a specific stimulus speed could increase spontaneous approach responses. Surprisingly, we found that introducing motion in this manner led to frequent freezing responses at all speeds of motion tested (**Fig. 2A**). Overall, from stimulus onset, mice froze more frequently and earlier (**Fig. 2A & B**). Approaches were most frequently observed when the stimulus was moving at the slowest presented speed of 2cm/s (**Fig. 2A-D**), while freezing was observed in the most trials when the stimuli moved the fastest, 50cm/s (**Fig. 2A-C and E**). Interestingly, after freezing mice often initiated an approach, and we observed that freezing responses preceded approaches 48% of the time. In addition, the proportion of approaches preceded by freezing steadily increased as stimulus speed increased, 31%, 63% vs. 80% at stimulus speeds of 2cm/s, 15 cm/s and 50 cm/s, respectively. Thus, these two behavioral responses to our stimuli were correlated, yet, oppositely biased by stimulus speed. We therefore calculated an approach/freeze behavior index which divided the number of approaches minus the number of freezes by the total number of observed behaviors of both types (**Fig. 2F**). This metric showed a clear relationship between the choice to approach or freeze in response to the specific speed of stimulus motion (**Fig. 2F**).

Importantly, our data also indicate that animals found each stimulus similarly behaviorally salient. The percentage of trials where animals exhibited at least one approach or freezing event was not significantly different as a function of stimulus speed (91%, 78% vs. 87% of 2cm/s, 15 cm/s vs. 50 cm/s conditions, respectively, N=23, Fisher’s exact test, p > 0.05 for all pairwise comparisons, **Fig. 2A**). Consistent with this, we found that the percentage of trials with a behavioral response of either type was significantly higher than those without either approaches or stimulus-driven freezing (N=23, number of trials with orienting responses, 21, 18 and 20, Fisher’s exact, p < .0001, p < 0.05 and p < 0.001, each speed respectively). Thus, the speed of the stimulus did not significantly preclude detection. Indeed, all stimulus speeds were behaviorally salient as they evoked at least one type of visual response with high probability, between 78-91%.

### Behavioral choice depends on relative stimulus size and speed as well as head-angle

Our assay provided the opportunity to estimate relative sizes and speeds of stimuli that best drove specific behavior as we could calculate the angular size and speed of stimuli as they would appear at the retina preceding specific behaviors. We hypothesized that the probability of approach would be inversely proportional to the size of the stimulus (mice preferring relatively smaller stimuli) and proportional to increasing speeds of motion (enhancing stimulus salience). However, the relationship was more complex. Regarding approach starts, mice were most likely to approach the slowest objective speed of stimuli. Therefore, stimuli were on average an angular size of approximately 5 deg of the visual arc as, on average, mice began their approaches to this stimulus from 24 cm away (**Fig. 3A, B&D**, green). Similarly, the preferred speed of a stimulus that preceded approach in mice was an average of 6 deg/s (**Fig. 3E**). On the other hand, distinct features drove freezing (**Fig. 3A – E**). Relatively smaller stimuli, roughly 4 deg in size and moving at a relative speed of greater than 80 deg/s biased behavioral choice towards freezing.

**Figure 3.**
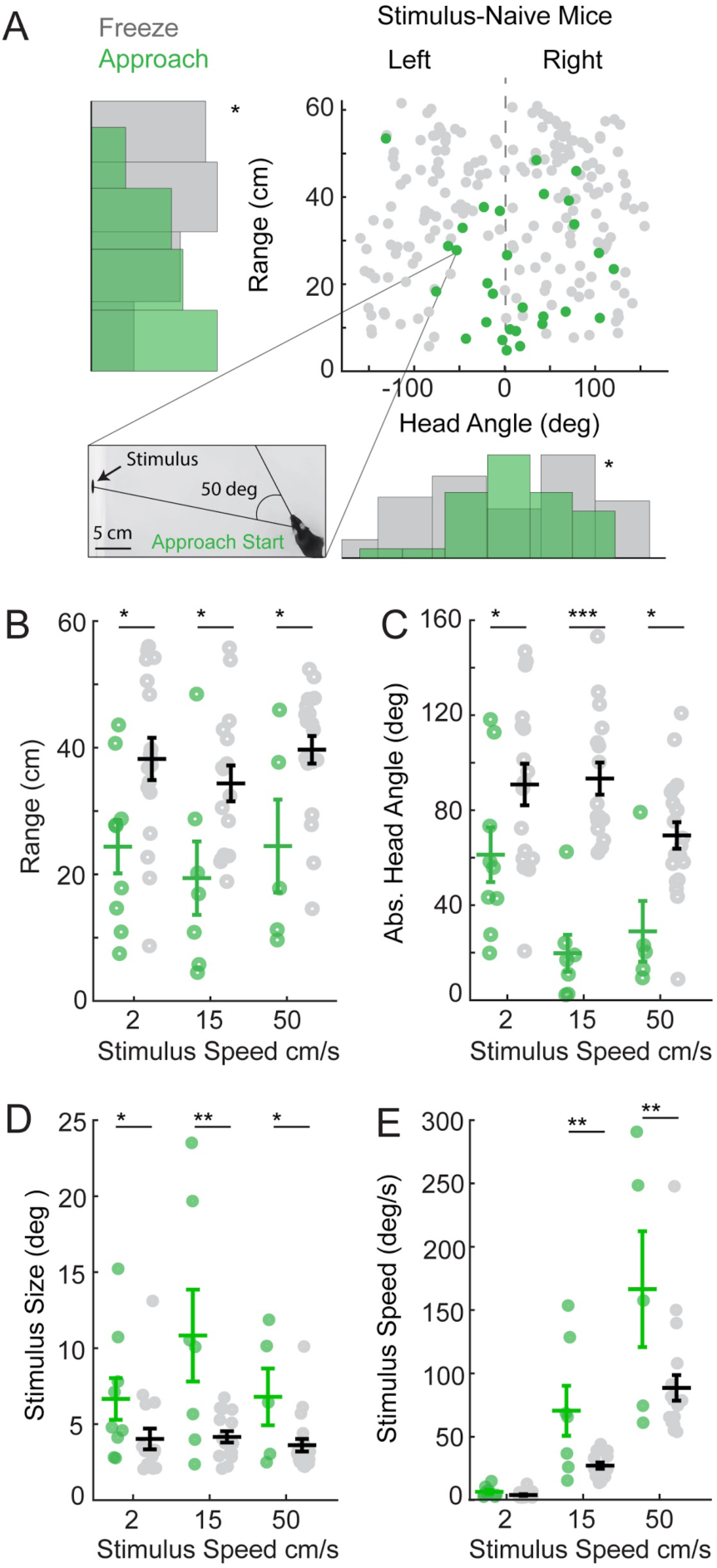
Freeze and approach orienting responses in naive mice are evoked by distinct combinations of stimulus size, speed, and location in the visual field. (A) Distribution of mouse ranges (cm) and head angles (deg) relative to stimulus location, where freezes occur (grey) or approaches start (green). Inset, representative frame from a video where the highlighted data point is measured. The frame is annotated to show the mouse’s head angle relative to the stimulus when that particular approach started. The range distributions for approaches versus freezes are significantly different from each other (KS test, * = p < 0.05) and show opposite skews towards far ranges for freezes and near ranges for approaches. The distribution of head angles between approach start positions and freeze positions are also significantly different (KS test, * = p < 0.05) and the distribution is unimodal for approach starts and bimodal for freezes (Ashman’s D > 2 for freezes). Plots show all individual behavioral events for all 23 mice, N = 29 and 199 approach starts and freezes, respectively. (B) Mean trial-averaged range at all three speeds where approaches started (green circles) or freezing occurred (grey circles) (two-way ANOVA, (F(1) = 19.99, p < 0.0001, Tukey’s post hoc, * = p < 0.05, N = 9, 7, 5 mice and 17, 16, 20 mice for each stimulus speed, approaches or freezes, respectively). (C) Mean trial-averaged absolute head angles of the mouse relative to the stimulus where approach starts or freezes were detected (two-way ANOVA, (F(2) = 3.17, p < 0.05, Tukey’s post hoc, * = p < 0.05, *** = p < 0.0001). (D) Mean trial-averaged estimated angular stimulus size where approach started or freezing was detected; calculation based on range and known objective size of the stimulus at its widest dimension (two-way ANOVA, (F(1) = 20.83, p < 0.0001, Tukey’s post hoc, * = p < 0.05 and ** = p < 0.01). (E) Mean trial-averaged estimated angular stimulus speed (relative speed) where approach started or freezing was detected (two-way ANOVA, (F(2) = 4.76, p < 0.05, Tukey’s post hoc, ** = p < 0.01). Error bars are +/-SEM.

Interestingly, mice also displayed clear differences in behavioral choice depending on where stimuli were likely detected within a predicted visual gaze location based on head angle measures (see methods for head angle calculations). Approaches were more likely to start when stimuli were located within a 65 deg head angle (**Fig. 3C**, green circles). In contrast, freezes were more likely to occur when moving stimuli were likely detected in the peripheral visual field, > 65 deg absolute head angle (**Fig. 3C**, grey circles). Notably, approaches could start from wider angles, but were unlikely, and freezes could occur with more centered head angle, but again less likely. As head angle predicts probable gaze direction, this has implications for which region of the visual field stimuli must appear in order to generate specific orienting behaviors (Michaiel, Abe, and Niell, n.d.). Importantly, these spatial biases are unlikely to be explained by biases in mouse preference for specific regions of the testing environment as stimuli were randomly presented on either screen and we observed no spatial bias in the arena in the absence of stimuli (16.7 ± 6.2%, 18.3 ± 4.9%, 18.3 ± 5.5%, 13.3 ± 4.4% percent time in each quadrant, 1-4, respectively during the baseline period, N=23, pairwise Wilcoxon rank sum tests, p > 0.05 for all pairwise comparisons). Stimuli thus appear randomly to either the right or left of the mouse and at variable ranges. Our data support the idea that distinct combinations of visual features consistent with distant moving objects relative to the subject selectively drive freezing behavior when perceived near the lower visual field. Furthermore, freezing may be linked to approach responses, yet are also dissociable based on the specific information that immediately precedes each type of behavioral response.

### Prey capture experience selectively increases approach frequency to specific stimulus speeds

Results presented so far were in mice that had not ever experienced hunting for prey and we predicted that the innate responses we observed related to prey detection and pursuit behaviors. As mice get more efficient at capturing live prey with experience (Hoy et al. 2016), we hypothesized that prey capture-experienced mice may increase their responsiveness to specific stimulus features that match real prey that they have successfully captured. To understand how the behavioral responses to motion stimuli observed in naïve mice related to prey capture-experienced mice, we quantified visually-evoked orienting responses in mice that had first experienced success in capturing crickets in the same environment (see Methods). First, we measured objective speed-dependent approach and freezing responses in prey capture-experienced mice (**Fig. 4**) in order to establish how this treatment affected relative stimulus size and speed-evoked behaviors (**Fig. 5**). We then directly compared the significant differences in preferences between naive and prey capture-experienced mice (**Fig. 6**).

**Figure 4.**
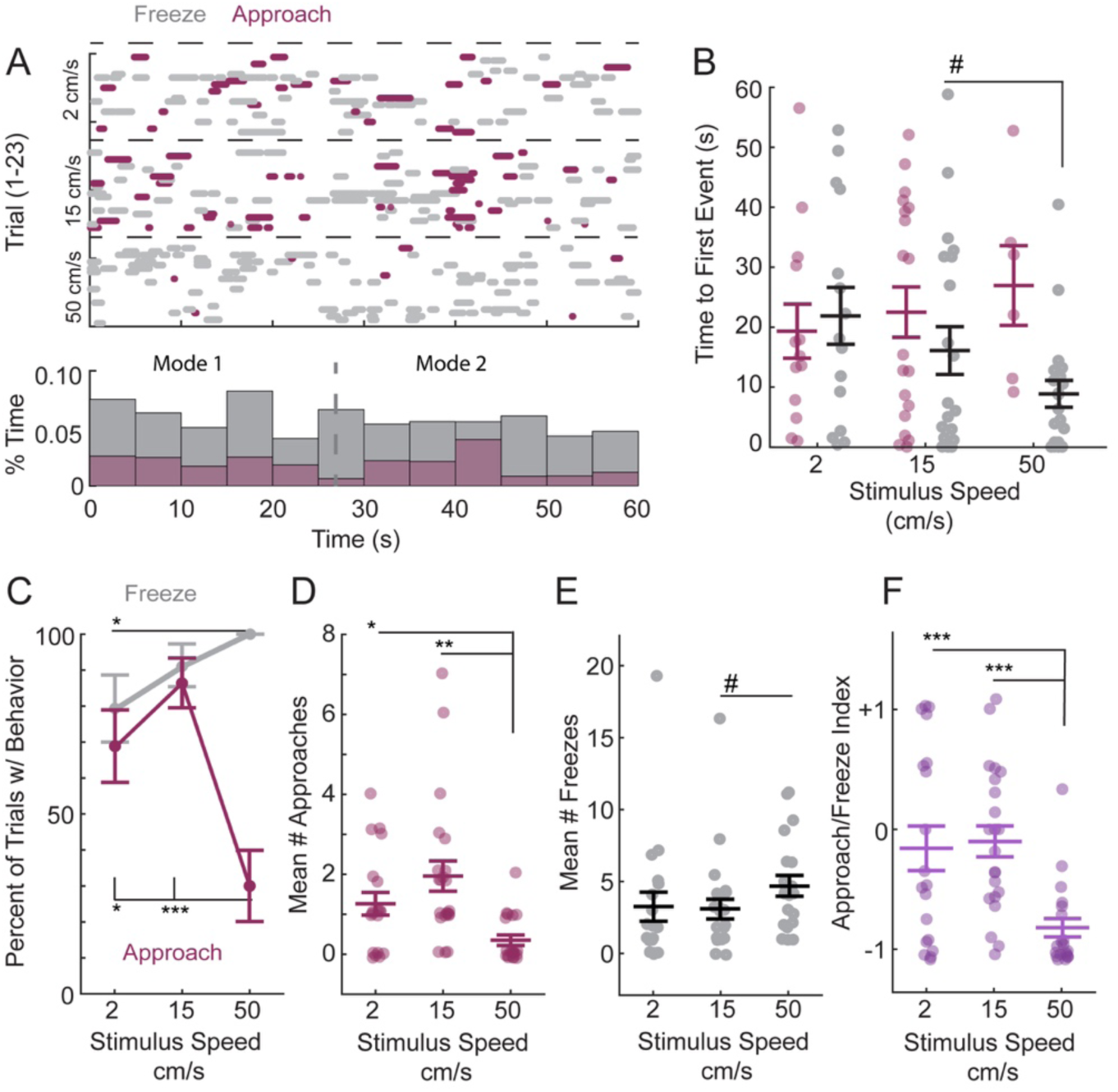
Mice with prey capture experience strongly prefer to approach moving stimuli with speeds less than 50 cm/s. (A) Top, an ethogram representing when approaches and freezes occurred from the onset of presentation of three different speeds of the stimulus for each mouse. Magenta = approaches by prey capture-experienced mice, grey = freezing displayed by prey capture-experienced mice. N = 23 mice performing one trial at each of three stimulus speeds, dashed line separates speed trial types. Bottom, histogram showing the proportion of time spent engaged in each type of response during each 5 s segment of the 60 s trial collapsed across all speeds. The distribution of approaches is bimodal and 26 s separates the modes (vertical grey dashed line). (B) Mean time to the first behavioral event of each type for mice that displayed at least one type of orienting response (two-way ANOVA, (F(1) = 9.1957, p < 0.06, Tukey’s post hoc, # = 0.0745, Ns = 13, 19, 6 mice and 15, 20, 20 mice for each speed, approaches versus freezes, respectively. (C) Percentage of trials at each speed with freezing (grey) or approach (magenta) showing significant differences in the observation of each type of behavior and that freezing probability steadily increased with objective stimulus speed (Fisher’s Exact test, * = p < 0.05 and **= p < 0.001 and adjusted r^2^ = 0.89). Error bars are standard deviation. (D) Number of approaches per mouse (one-way ANOVA, (F(2) = 9.68, p < 0.001, Tukey’s post hoc, * = p < 0.05 and *** = p < 0.001). (E) Number of freezes per mouse. The effect of stimulus speed on the number of freezes observed during a trial between speeds approached significance (# = p=0.072). (F) Approach-to-freeze index calculated as shown in Figure 2 (one-way ANOVA, (F(2) = 13.11, ***p < 0.0001, Tukey’s post hoc, *** = p < 0.0001). Error bars are +/-SEM.

**Figure 5.**
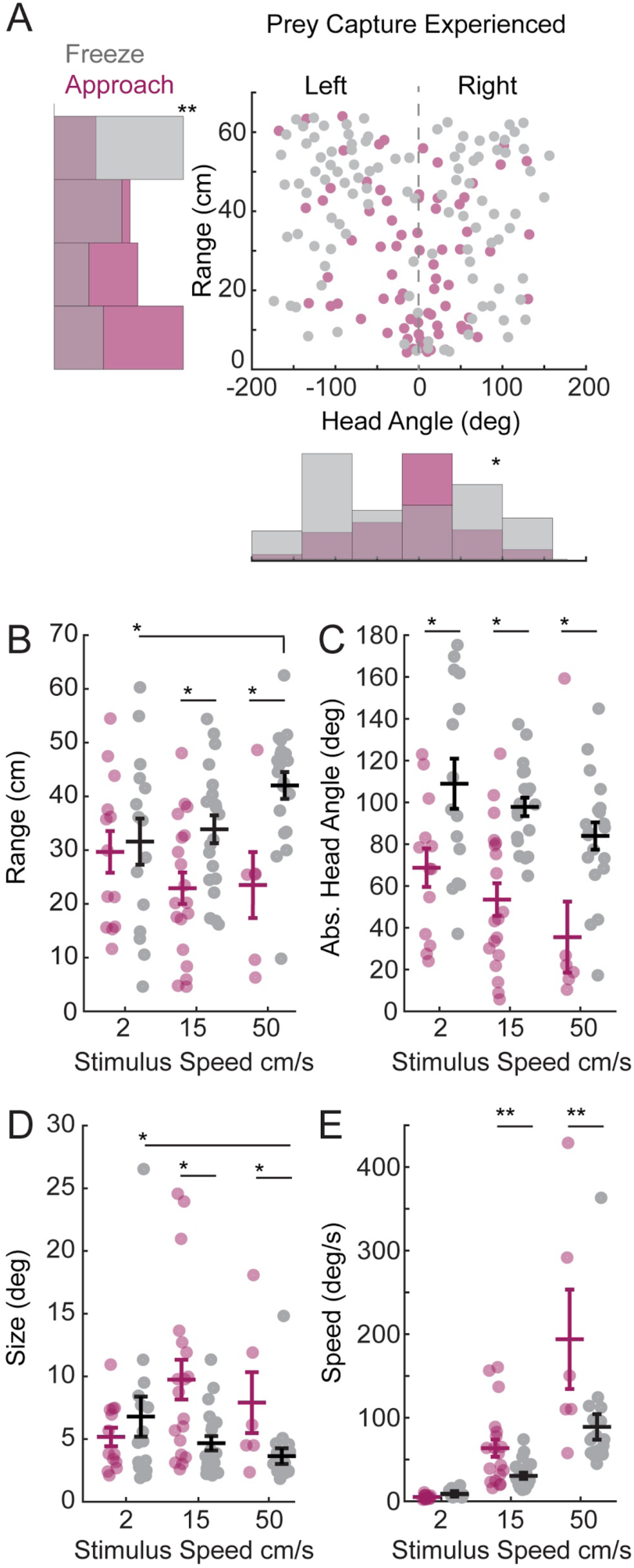
Prey capture-experienced mice prefer to approach objects with distinct visual features from those that drive freezing. (A) Distribution of mouse ranges (cm) and head angles (deg) relative to stimulus location where freezes occur (grey) or approaches start (magenta). The range and head angle distributions for approaches versus freezes are shifted relative to each other, showing opposite skews towards far ranges for freezes and near ranges for approaches (KS test, p < 0.01). We observe a bimodal distribution of head angles towards the periphery for freezes, and normal distribution well-centered near 0 deg for approaches. All individual behavioral events for all 23 mice are plotted (N = 74 and 224, approaches versus freezes, respectively). (B) Mean trial-averaged range at all three speeds where approaches started (magenta circles), or freezing occured (grey circles) (two-way ANOVA, (F(1) = 12.49, p < 0.001, Tukey’s post hoc, * = p < 0.05, Ns = 13, 19, 6 mice and 15, 20, 20 mice for each speed of stimulus, approach versus freezing data, respectively). (C) Mean trial-averaged, absolute head angles of the mouse relative to the stimulus where approach started or freezing occured showing that approaches were initiated at more direct head angles than freezes, and this separation increased as stimulus speed increased.(two-way ANOVA, (F(1) = 44.39, p < 0.0001, Tukey’s post hoc, * = p < 0.05) (D) Mean trial-averaged estimated angular stimulus size where approaches started or freezing occured, based on range and known objective size of the stimulus at its widest dimension. Stimuli that evoked approach or freeze depended on stimulus speed (two-way ANOVA, (F(2) = 4.89, p < 0.05., Tukey’s post hoc, * = p < 0.05). (E) Mean trial-averaged estimated angular stimulus speed where approach starts or freezes were detected (two-way ANOVA, (F(2) = 5.91, p < 0.01, Tukey’s post hoc, ** = p < 0.01). Error bars, +/-SEM.

**Figure 6.**
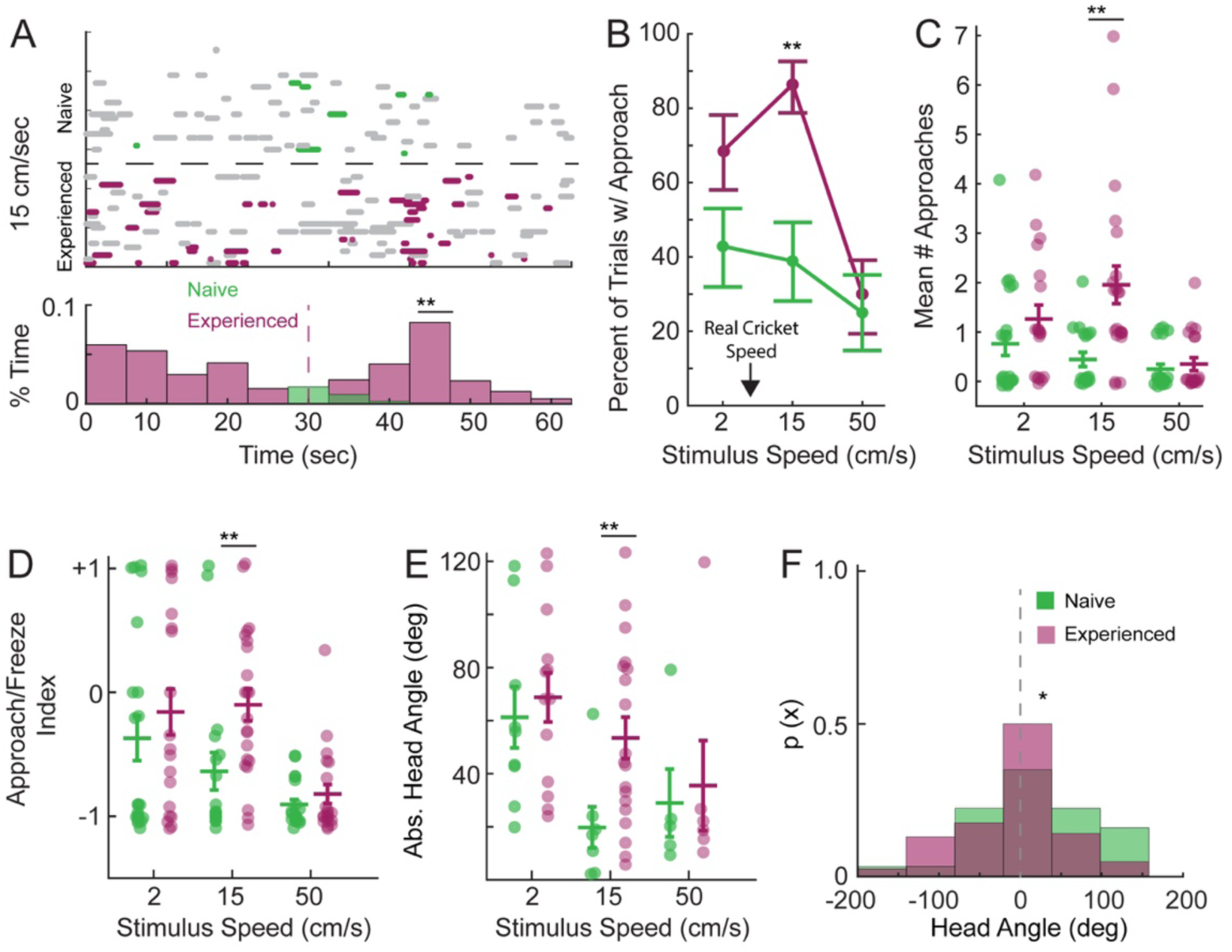
Prey capture experience selectively alters approach frequency towards specific objective speeds of stimuli and where stimuli are detected in the visual field. (A) Top, direct comparison of previously shown ethograms representing when approaches started and freezing occurred from the onset of the presentation of the 15 cm/s speed stimulus for each mouse, previously shown in **Figs. 2A and 4A**. Magenta = approaches by prey capture-experienced mice, green = approaches by naive mice. Grey = freezes in either naive (top panel) or prey capture-experienced (bottom panel) mice. The overlaid histograms of these distributions reveals that prey capture-experienced mice respond earlier in the trial and more frequently to the 15 cm/s speed stimulus relative to naive mice (KS test, ** = p < 0.01, N =8 vs. 43 approach events from naive vs. prey capture-experienced mice, respectively, from the 15 cm/s condition). Effects at other objective speeds are insignificant, data not shown together. (B) Percent of trials where approaches were observed for all three speeds of stimuli revealing a highly significant difference between naive and prey capture-experienced mice for the 15 cm/s stimuli (Fisher’s Exact test, ** = p < 0.01, N=23). Error bars are standard deviation. Black arrow highlights average measured crawling speed of crickets. (C) Mean number of approaches per mouse compared directly between naive (green) and prey capture-experienced (magenta) mice reveals that the frequency of approach selectively increases for the 15 cm/s speed stimulus (Tukey’s post hoc, ** = p < 0.01, N= 21, 18 and 20 for naive mice and N = 19, 22 and 20 for prey capture-experienced mice, for each speed, respectively). (D) Approach/freeze index showing that the selective increase in approach frequency evoked by the 15 cm/s stimulus in prey capture-experienced mice leads to a significant increase in the ratio of the two types of orienting responses at only this objective stimulus speed (Tukey’s post hoc, ** = p < 0.01). (E) Mean trial-averaged absolute head angle of approach starts directly compared between naive (green) and prey capture-experienced (magenta) mice (Tukey’s post hoc, ** = p < 0.01). (F) The distribution of all head angles at approach start for all speed for naive (green) versus prey capture-experienced (magenta) mice, reveals a shift in the visual field location where all stimuli are detected prior to approach (KS test, * = p < 0.05, N= 29 and 73 approaches for naive vs. prey capture-experienced mice, respectively). Error bars are +/-SEM.

Notably, prey capture-experienced mice quickly (within 25 s) and selectively approached stimuli with objective speeds of 2 cm/s and 15 cm/s speeds but not the 50 cm/s stimulus (**Fig. 4A & B**, 39% vs. 43% vs.13%, of mice approached within 25 s, each speed respectively from slow to fast, N= 23, Fisher’s exact test, p < 0.05, 2 cm/s vs. 50 cm/s and p < 0.05, 15 cm/s vs. 50 cm/s, respectively). Instead, mice immediately and selectively froze in response to the 50 cm/s stimulus (**Fig. 4A & B**). Similarly, the percentage of trials with an approach overall was significantly higher for the 2 cm/s and 15 cm/s speed stimulus versus the 50 cm/s stimulus, while the percentage of trials with freezing steadily increased as stimulus speed increased (**Fig. 4C**). These selective increases in approach behaviors towards the 2 and 15 cm/s stimulus speeds coupled to reduced freezing responses relative to the 50 cm/s stimulus, were further supported by observing a highly significant increase in the approach to freeze index for the slower speeds of stimuli (2 and 15 cm/s) versus the fastest (50 cm/s; **Fig. 4D-F**). Altogether, these data demonstrate that prey capture-experienced mice are highly selective for the stimuli that induce approach versus freezing and stimuli are detected early within the trial.

To understand how prey capture experience may impact the relative features of visual stimuli that drive approach responses, we estimated the angular size and speed of the stimulus as well as relative head angle when prey capture-experienced mice began approaches or began freezing. Approaches started towards stimuli ranging from 5-10 deg and moving between 5-60 deg/s and when stimuli likely appeared in the central region of the predicted visual gaze (**Fig. 5**, magenta). In contrast, freezing was initiated by 4 deg stimuli moving at greater than 80 deg/s and localized to the periphery (**Fig. 5A**). Intriguingly, these findings demonstrated that prey capture experience selectively increased approach responses to selective visual features of stimuli relative to mice naive to prey capture. We therefore directly compared key measures of approach behaviors between prey capture-naïve and -experienced mice (**Fig. 6**). First, prey capture experience significantly increased the proportion of stimulus presentation trials where approaches were observed when stimuli moved objectively at 15 cm/s (**Fig. 6A-C**). This corresponded to a highly significant increase in approach frequency and the approach/freeze index when mice were presented the 15 cm/s stimulus (**Fig. 6C & D**). These findings suggested that prey capture experience selectively altered the innate orienting responses evoked by our novel stimuli nearer the range of speeds exhibited by live prey, less than 15 cm/s. The average speed of crawling locomotion for crickets, the prey used in this study, was 5 +/-2.3 cm/s and was closest to the slowest objective speed of stimulus that we tested that innately drew the most approaches in naive mice (**Fig. 2D**).

Surprisingly, the most striking evidence that prey capture experience altered visual processing of stimuli is that mice were more likely to approach the 15 cm/s stimulus (**Fig. 6B & C**) and detect it earlier in the trials than naïve mice as indicated by the more frequent observation of approaches starting before 25 s (**Fig. 6A**). Additionally, mice were significantly more likely to initiate approaches when stimuli appeared at peripheral head angles greater than 60 deg (**Fig. 6E**). Interestingly, this is the average head angle preceding approaches to the slowest speed of stimuli that naïve mice most strongly prefer to approach (**Fig. 6E**). Furthermore, though we noted that more approaches could start from peripheral head angles in prey capture-experienced mice for specific speeds of stimuli, head angles were less variable overall around the center of the head angle distribution which predicts probable visual field location (center of head angle distribution = +10 deg and +10 deg, width at half max = 180 deg vs. 60 degrees, naive vs. prey capture-experienced mice, respectively) suggesting that stimuli were detected more equally between the predicted hemifields of the overall visual field (**Fig. 6F**). Together, these data suggest that prey capture experience, and therefore possibly the expectation of prey, selectively altered the features of visual stimuli that evoke innate approach responses towards novel stimuli and enhanced stimulus salience.

Finally, given the observation of frequent freezing responses, we sought to explore further whether these responses more likely reflected behaviors relevant to prey capture versus threat detection. We therefore quantified the probability that mice responded to our stimuli with additional appetitive behaviors, interception and pursuit (followed after an initial approach), or active avoidance behavior such as spending more time near the periphery of the arena and corners (thigmotaxis). We reasoned that if the freezing indicated threat detection, we should see an increase in additional avoidance behaviors as objective stimulus speed increases as that is when the freezing responses are highest. We also predicted that if our fastest moving stimuli were perceived as threatening, we should not observe significant appetitive responses to these stimuli. However, we found that both naive and prey capture-experienced mice significantly intercepted and pursued all speeds of our moving stimuli above a hypothetical mean of 0 (**Fig 7A-C, Videos 1, 2, 4 & 5**) and displayed similar levels of thigmotaxic behavior relative to baseline conditions without stimuli (79 ± 5.6 % thigmotaxis in baseline condition without stimuli and Hoy et al., 2019). Thus, overall mice responded with significant levels of appetitive type responses to all stimuli and did not modulate overt stimulus avoidance behaviors (Hoy et al. 2016; Blanchard, Flannelly, and Blanchard 1986). However, their rate of interception given an approach start and probability of pursuit varied by stimulus speed and experience (**Fig. 7B & C**). Therefore several distinct appetitive behaviors are significantly modulated by, and therefore relevant to, prey capture experience. Overall, these findings suggest that despite a high frequency of freezing responses to the presented stimuli in both naïve and prey capture-experienced mice, mice did not display classically defined overt signs of anxiety or active avoidance of our stimuli. Instead, mice did display clear signs that all stimuli could be perceived as appetitive yet also clearly revealed strong preferences to release appetitive behaviors towards specific features. Taken together, these observations suggest that prey capture experience selectively increases appetitive, stimulus pursuit behaviors towards specific features of visual stimuli and that freezing is unlikely to indicate threat in this context.

**Figure 7.**
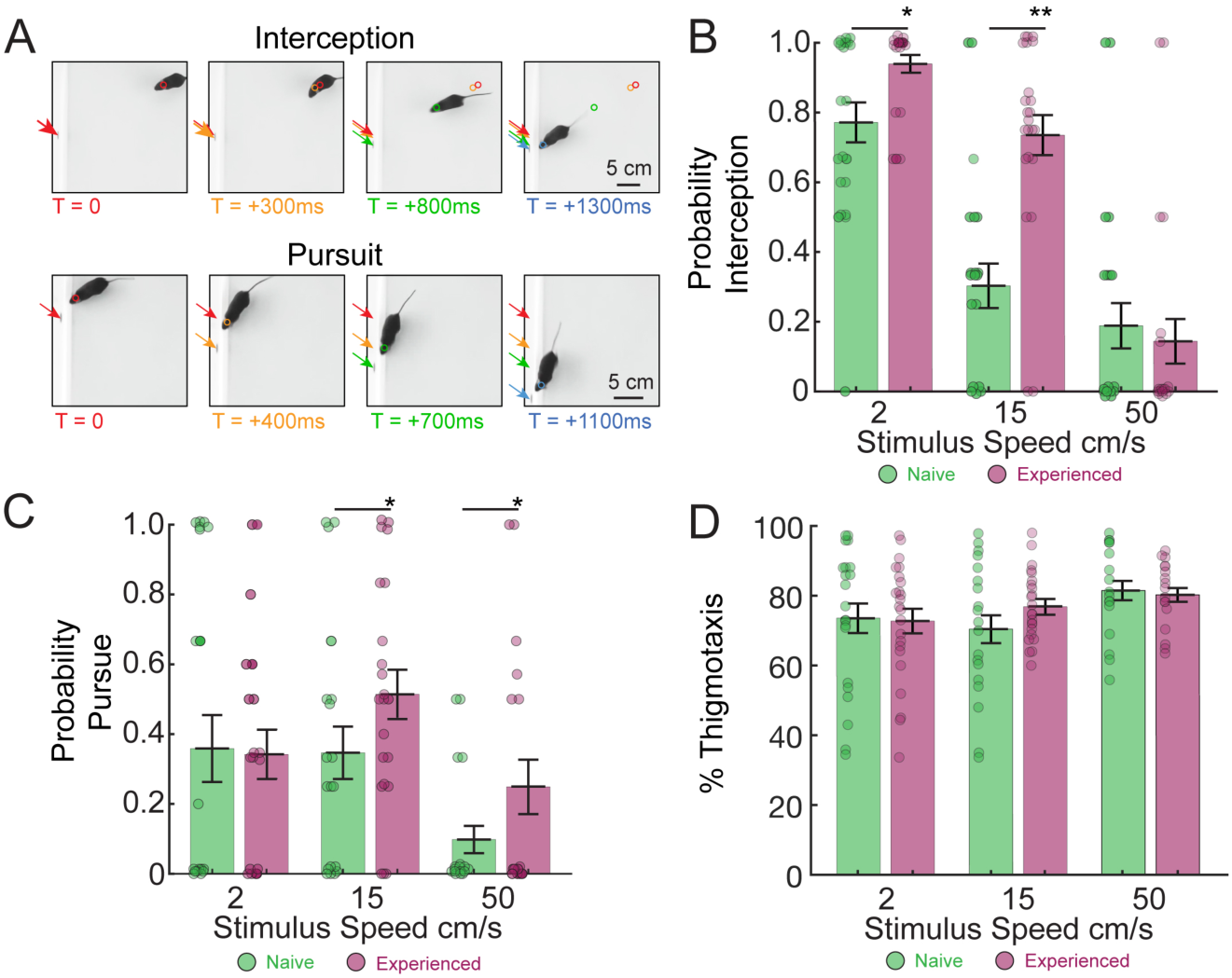
Mice perceive moving stimuli as appetitive and prey capture experience significantly enhances specific appetitive behaviors towards specific objective speeds. (A) Representative appetitive behaviors scored by three human observers with an inter-scorer reliability of > 95% (Cohen’s K = 0.90): stimulus interception (top) and stimulus pursuit (bottom). Colored arrows indicate the position of the stimulus in sequential moments in time identified by the colored text below. Colored circles indicate head position at the same times highlighted by each color. (B) Mean trial-averaged interception probability across mice that displayed at least one approach start when presented each speed of stimulus. (C) Mean trial-averaged pursuit probability across mice that displayed at least one approach start when presented each speed of stimulus. (D) Mean trial averaged percent time spent within 10 cm from the periphery of the arena (thigmotaxis) across mice that displayed at least one approach start. In all panels, Student’s t test, pairwise comparison between naive vs. prey capture-experienced mice, * = p < 0.05, ** = p < 0.01, N = 21, 19 & 17 and 23, 23 and 17 mice at each speed, naive vs. prey capture-experienced, respectively. Error bars, +/-SEM.

## Discussion

Our observations reveal an important and novel understanding of the specific visual features that innately evoke approach versus freezing responses in C57BL/6J mice. Overall, naive mice preferentially approached a stimulus that spanned 5 deg of the visual field, moved at 6 deg/s and was detected within a 65 deg head angle relative to the stimulus. These stimulus preferences that evoked rapid approach are consistent with previous studies of measured approach responses towards live crickets during prey capture (Hoy et al. 2016). On the other hand, mice preferentially froze in response to a 4 deg stimulus that moved faster than approximately 80 deg/s and was detected at head angles greater than 65 deg. Thus, our observations demonstrate that specific combinations of size, speed and visual field location of novel stimuli enhance stimulus detection and evoke distinct orienting responses.

While others have previously noted that specific stimulus features may cause freezing when presented overhead (Yilmaz and Meister 2013; De Franceschi et al. 2016), our data demonstrate that selective visual features evoke freezing when perceived near the horizon of the environment of the mouse (**Fig. 2-5**). Indeed, two recent studies precisely quantifying eye movement as coupled to head movement in mice engaged in two different ethological contexts including prey capture showed that the position of the eyes, and therefore probable visual field location and predicted gaze, track the head position along the azimuth and in elevation (Michaiel, Abe, and Niell, n.d.; Meyer, O’Keefe, and Poort, n.d.).In addition, we quantified behavioral responses that are traditionally associated with either appetitive versus aversive conditions, specifically comparing interception and pursuit to thigmotaxis behavior (**Fig. 7**). This analysis further supports our interpretation that freezing responses in this context most likely reflect a distinct perceptual process from those measured in “looming” assays, as we often observed appetitive responses to all of our moving stimuli and failed to observe overt aversion in the form of increased thigmotaxis.

Freezing behavior is hypothesized to facilitate visual processing in nonthreatening contexts where ceasing self-motion upon the detection of suddenly appearing stimuli can help with accurately perceiving external motion (Roseberry and Kreitzer 2017; Roelofs 2017) and augment the perception of specific visual features (Lojowska et al. 2015). Consistent with this idea, we found that at least one freeze preceded approaches 48% of the time in our naive mice (**Fig. 2A & B and 5A & B, Video 1 & 4**) and mice followed the trajectory of moving stimuli with saccadic head movements while retaining a frozen body posture (**Video 4 & 6**). However, future studies that apply recently improved dimensionality reduction methods to the complex behaviors and behavioral sequences that we observe here will further advance our understanding of how freezing differs depending on context (Berman, Bialek, and Shaevitz 2016; Datta 2019; Datta et al. 2019). Furthermore, how freezing in response to specific stimuli relates to specific physiological states and perceptions may be partially determined via simultaneous measure of parasympathetic versus sympathetic nervous system activation (Roelofs 2017). It will therefore be interesting in future experiments to record the neural activity of identified cell types that encode the stimulus features shown to drive freezing here in conjunction with measures of autonomic nervous system activation under our experimental conditions as compared to those induced by the presentation of overhead looming stimuli that also cause freezing responses.

Altogether our study will aid in interpreting both past neural circuit studies in mice as well as facilitate the improvement of future studies seeking to decode visual perception and visuomotor transformations in the mouse model. For example, we found that not all relatively small moving novel stimuli in the lower visual field reflexively trigger approach (Dean, Redgrave, and Westby 1989; Shang et al. 2019). Indeed, overt appetitive response choices in the naive, freely moving animal were clearly evoked more readily by specific combinations of features. In contrast, when the possibility of a prey reward was altered via natural prey capture experience prior to stimulus exposure, we saw a selective modification of approach response evoked by distinct aspects of visual information. Thus, our work provides a roadmap to the specific types of experiences to manipulate and the range of visual stimulus features that should be monitored in future studies of the mouse visual system attempting to determine the neural activity that encodes information behaviorally-salient to the mouse. Our findings also further bolster the notion that neural circuits mediating different types of visual orienting responses may be dissociable by their specific sensory inputs. Furthermore, these inputs would encode a specific combination of stimulus size, speed of motion, and position in the visual field. In the mouse model, we are coming to appreciate the richness of visual features computed in the early visual system, including at the level of the retina (Farrow and Masland 2011; Yonehara and Roska 2013; Jacoby and Schwartz 2017). It will therefore be interesting in future studies to determine whether specific information first computed in the retina and relayed via distinct retinal ganglion cell types to different projection targets is sufficient to rapidly drive each type of orienting response, or whether it must be further transformed or filtered in target regions in order to generate observed behavioral responses (El-Danaf and Huberman 2019; Lee et al. 2020). Almost ninety percent of RGCs in the mouse project to the superior colliculus, which contains cells with response properties similar to those found in the retina (Gale & Murphy, 2014) and is known to mediate visual attention (Krauzlis et al. 2013; Knudsen 2020) and orienting (Dean & Redgrave, 1984; Overton et al., 1985; Westby et al., 1990). In the zebrafish, discrete RGC types have indeed been shown to underlie approach, while visual stimulus feature-driven decisions to approach or avoid are computed and encoded in the optic tectum, homologous to the superior colliculus (Semmelhack et al. 2014). Interesting future studies will seek to address whether similar circuit mechanisms exist in the retino-collicular pathways of the mouse that underlie the orienting response choices measured in our current study (Procacci and Hoy 2019; Shang et al. 2019; Zhao et al. 2019).

Excitingly, our study also reveals flexibility as well as specificity in how mice respond to specific visual cues naturally, this will pave the way for identifying the neural mechanisms that underlie visual processing of specific appetitive stimulus features and how their representations may change as a function of additional sensory cues, behavioral state, development and experience. Therefore, perhaps the most noteworthy aspect of the current study is that we quantitatively demonstrate that mice preferentially approach novel objects of specific size, speed, and location in the visual field, and that the salient features of objects that evoke approach are refined selectively by prey capture experience. The findings that mice are more likely to approach a specific stimulus objectively traveling between 2-15 cm/s relative to naïve mice (**Fig. 6A-D**) and detect it both earlier in the trial (**Fig. 6A**) and more in their periphery than naïve mice (**Fig. 6E**), argue that prey capture experience modifies specific aspects of visual processing. A distinct possibility was that a rewarding experience, such as successful prey capture, might more generally increase motivation to approach all stimuli, or reduce freezing responses to all stimuli. However, this did not occur. Instead, we found that freezing probability did not change significantly after a rewarding experience, nor did the features of visual stimuli that evoked freezing responses change. Our data demonstrate that prey capture experience and/or the expectation of prey in the environment selectively alters the activity of visual pathways that transform specific visual information into approach and pursuit behavior in the prey capture context. Therefore, one of the more exciting avenues of further research stemming from this work, is to determine the underlying circuit mechanisms driving conditional approach responses to specific stimuli in the mammalian brain. This research direction may ultimately enhance our understanding of the nature of exploratory behavior and learning within natural, appetitive contexts.

## Materials and Methods

### Subjects

All experiments were conducted in accordance with protocols approved by the University of Nevada, Reno, Institutional Animal Care and Use Committee, in compliance with the National Institutes of Health Guide for the Care and Use of Laboratory Animals. Ten C57BL/6J mice between the ages of 2 and 4 months were used for experiments in which a stationary stimulus was varied in size. Twenty-three C57BL/6J mice between the ages of 2 and 4 months were used in each of two groups, naïve and prey capture-experienced, for experiments in which stimulus speed was varied. Mice were group-housed, up to 5 animals per cage, with regular access to water and food (Envigo, Teklad diet, 2919) in an on-campus vivarium. The vivarium was maintained on a 12 hour light/dark schedule, and all testing occurred within 3 hours of the dark to light transition. The crickets used for prey capture were *Acheta domestica*, 1-2 cm in length, obtained from Fluker’s Farm. They were group-housed in a cage and fed Fluker’s Orange Cube Cricket Diet.

### Visual Stimuli

Visual stimuli were generated with MATLAB Psychophysics toolbox (Brainard 1997) and displayed on LCD monitors (60 Hz refresh rate, ∼50 cd/m^2^ luminance) in a darkened room. The computer monitors replaced two sides of a rectangular behavioral arena that was 60 cm × 60 cm × 30 cm, length x width x height. Stationary visual stimuli were centered between 2-15 cm from the bottom of the arena depending on their size. To mimic probable insect proportions, we displayed ellipses with a major axis (width) that was 2 times the size of the minor axis (height), and we varied the major axis from 0.5 to 10 cm. This approximates a low pass filtered image of a cricket (shape lacking detailed texture) viewable at the horizon between floor and wall. For a stimulus with a major axis of 2 cm, this corresponds to the presentation of an approximately 6 deg target from 20 cm away from the screen. 2 × 1 cm ellipses evoked the most approaches from naive mice with the highest pursuit speeds (pursuit speed by stimulus size); we therefore kept this size constant when we varied stimulus speed for a separate cohort of animals. Objective stimulus speed was varied over 3 steps between 2 cm/s to 50 cm/s and presented in a random order to each mouse. Mice could therefore view a stimulus moving at speeds varying from 2 to ∼300 deg/s at the retina. This is consistent with a range of speeds presented to ferrets (Ritter, Anderson, and Van Hooser 2017) and the range of speed selective responses that are encoded in the superior colliculus of mice (Gale and Murphy 2018, 2014; Inayat et al. 2015; Wang et al. 2010).

### Behavior

Prior to testing, mice were acclimated to handlers for 2 days during which each mouse was handled 3 times a day for 5 min each. Mice were then acclimated to the arena for up to 4 days during which each mouse was placed in the arena 3 times a day for 5 min each. The arena floor was cleaned thoroughly with 70% EtOH after each mouse was removed to mitigate odor distractions.

On testing day, each mouse was placed in the arena for 3 min to habituate. The final 60 s of this period was used as a control trial to assess mouse behavior in the absence of a stimulus. After this period, a randomized order of stimulus speeds was presented to the mouse for 60 s each speed and an inter-stimulus-interval of 60 s.

For prey capture-experienced mice, each mouse was given a live cricket starting on their second day of habituation in the arena. If the mouse caught the cricket within the 5 min period of time, it was given another until it caught up to three over the 5 min period. If it did not catch the cricket, it was still returned to its home cage at the end of the session. All mice were then returned to their home cages with standard food only. This modified habituation protocol was repeated for up to 4 days to be comparable to the “naive” mouse group in terms of time spent in the environment; most mice exposed to crickets reliably caught crickets in the arena in less than 30 s after 3 days (>80%). These mice were then exposed to our virtual motion stimuli as described above.

### Data Analysis

DeepLabCut software (Mathis et al. 2018) was used to digitize and extract two-dimensional coordinates of the mouse nose, ears, and tail base, as well as the center point of the stimulus, throughout the duration of each trial. Tracks were entered into customized MATLAB scripts to extract behavioral parameters (Hoy et al., 2019). We wrote a custom function to identify freezes, defined as any time the mouse’s body center moved less than 0.5 cm/s for a duration of 0.5 – 5 s. Since mice often froze in the corners of the arena in the absence of a stimulus, corner freezes were excluded from our analysis of stimulus-evoked freezes. A successful approach in Figures 2-6 was defined as any time the mouse’s nose eventually came within 2 cm of the stimulus center after moving toward the stimulus for a distance of at least 5 cm, or at an average speed of at least 15 cm/s, in order to generate a conservative estimate of approaches and estimate stimulus features correlated to start of successful approaches.

Using these behavioral definitions, we computed the percentage of 2, 15, and 50 cm/s stimulus trials in which each type of response was observed, as well as the number of freezes and approaches that occurred during individual trials. Behavioral index scores were calculated by taking the number of approaches minus the number of freezes, and dividing by the number of approaches plus the number of freezes for each trial. For both response types, we calculated range as the distance between the central point between the two ears and the center of the stimulus, and bearing as the angular distance between a line emanating from the center point between the mouse ears to the mouse nose, and the line emanating from the center point between the mouse ears to the center of the stimulus. The center point between the mouse ears constituted the vertex. Angular stimulus sizes and speeds were calculated using the long axis (horizontal with respect to the ground, width) of the stimulus (2 cm) and the distance of the head from the stimulus at the moment we observed the relevant orienting behavior. Three human observers scored stimulus interception and pursuit (following after approach) behaviors. Their inter-scorer reliability was measured as Cohen’s K and based on the percent of events they agreed to include and those excluded. To gain conservative estimates of appetitive behavior we only considered those behaviors that had reliable score between observers. For example, interception was consistently defined as nose position coming within 2 cm of the stimulus at the end of an approach trajectory and generated a robust interscorer reliability, Cohen’s K > 95%.

### Statistics

Statistics were performed using MATLAB and R packages. Where means are reported, data were fitted to a linear mixed effects model in which we entered a combination of response type (freeze, approach), stimulus speed (2, 15, 50 cm/s) and/or experience (naïve versus prey capture-experience) as fixed variables, and individual subjects as random variables. This method accounted for repeated measures while also retaining data from animals with missing data points. We then performed one- or two-way ANOVAs, as indicated in figure legends, F(1) indicting main effect of stimulus speed, F(2) main effect of behavioral response type. Tukey’s HSD post hoc testing was used to determine whether two means were significantly different for specific groups and conditions. For single sample tests and a priori hypothesis testing, Student’s t-test then corrected for multiple comparisons was used to determine whether a particular group was significantly different than a hypothetical mean of 0 or from a specific experimental group predicted to be distinct. Where percentages are compared, Fisher’s Exact tests were used to determine significance. Tests were corrected for multiple comparisons. Significant differences in distributions of head angles, ranges or stimulus features between groups were detected with Kolmogorov-Smirnov tests and circular measures were confirmed with Kuiper’s test which is sensitive to differences in the tails of the distributions and multimodal distributions. We also employed Hartigan’s dip tests or Ashman’s D measures to determine whether data distributions were multimodal, Ashman’s D > 2 indicates two separable modes.

## Supporting information

Pose estimation

Stimulus induced Freeze

Complex stimulus induced behavioral sequence

pursuit behavior

Complex Freeze with re-orienting behavior

## Acknowledgements

We would like to thank Makayla Kesler-DeBarge and Rocio Olvera for their assistance in scoring behavioral responses in mice, running pose estimation routines and handling experimental mice. We thank Drs. Stephen Van Hooser, Cristopher Niell, Matt Smear, Angie Michaiel, Philip Parker and Thomas Kidd, as well as members of the Hoy lab, for reading manuscript drafts and providing helpful comments on text and data analysis. This work was supported by P20GM103650 (JLH) and partially supported by NSF Grant for an REU site in Biomimetics and Soft Robotics (BioSoRo) with award number EEC #1852578NSF (JLH and Dr. Yantao Shen).

## Author Contributions

Conceptualization, J.L.H.; Methodology and Formal Analysis J.L.H., N.P., K.A., and E. R.; Software, J.L.H., N.P., E. R.; Investigation, J.L.H., N.P., R. I., K.A.; Visualization, J.L.H. and N.P.; Writing – Original Draft, N.P.; Writing – Review & Editing, J.L.H., N.P., and K.A..; Resources, J.L.H.

